# RGS14 modulates locomotor behavior and ERK signaling induced by environmental novelty and cocaine within discrete limbic structures

**DOI:** 10.1101/2021.01.21.427672

**Authors:** Stephanie L. Foster, Daniel J. Lustberg, Nicholas H. Harbin, Sara N. Bramlett, John R. Hepler, David Weinshenker

**Affiliations:** Emory University School of Medicine, Department of Human Genetics, Atlanta, GA 30322; Emory University School of Medicine, Department of Pharmacology and Chemical Biology, Atlanta, GA 30322

**Keywords:** RGS14, central amygdala, hippocampal area CA1, hippocampal area CA2, novelty-induced locomotion, psychostimulant-induced locomotion, cocaine, c-fos, ERK, thigmotaxis

## Abstract

**Rationale:** In rodents, exposure to novel environments or psychostimulants promotes locomotor activity. Indeed, locomotor reactivity to novelty strongly predicts behavioral responses to psychostimulants in animal models of addiction. RGS14 is a plasticity restricting protein with unique functional domains that enable it to suppress ERK-dependent signaling as well as regulate G protein activity. Although recent studies show that RGS14 is expressed in multiple limbic regions implicated in psychostimulant- and novelty-induced hyperlocomotion, its function has been studied almost entirely in the context of hippocampal physiology and hippocampusdependent behaviors.

**Objective:** We sought to determine whether RGS14 modulates novelty- and psychostimulant-induced locomotion and neuronal activity.

**Methods:** We assessed *Rgs14* knockout (RGS14 KO) mice and wild-type (WT) littermate controls using novelty-induced locomotion (NIL) and cocaine-induced locomotion (CIL) behavioral tests with subsequent quantification of c-fos and phosphorylated ERK (pERK) induction in limbic regions that express RGS14.

**Results:** Compared to WT controls, RGS14 KO mice exhibited attenuated locomotor responses in the NIL test, driven by avoidance of the center of the novel environment. By contrast, RGS14 KO mice demonstrated augmented peripheral locomotion in the CIL test conducted in either a familiar or novel environment. The absence of RGS14 enhanced induction of c-fos and pERK in the central amygdala and hippocampus (areas CA1 and CA2) when cocaine was administered in a novel environment.

**Conclusions:** RGS14 regulates novelty- and psychostimulant-induced hyperlocomotion, particularly with respect to thigmotaxis. Further, our findings suggest RGS14 may reduce neuronal activity in discrete limbic subregions by inhibiting ERK-dependent signaling and transcription.

## Introduction

Locomotor activity in rodents is augmented by exposure to novel environments or psychostimulants, like amphetamine or cocaine, via enhanced signaling of the catecholamine neurotransmitters dopamine (DA) and norepinephrine (NE) within limbic circuits that command memory, motivation, emotion, and stress reactivity (Carmen Arenas et al. 2016; Fink and Smith 1980; Walker et al. 2009; Wingo et al. 2016). Novelty-induced locomotion (NIL) strongly predicts individual differences in sensitivity to psychostimulant effects in animal models of drug addiction (Arenas et al. 2014; Belin et al. 2011; Hooks et al. 1991; Vidal-Infer et al. 2012), suggesting that partially overlapping neurobiological substrates organize these behaviors (Badiani et al. 1998; Hooks and Kalivas 1995; Kabbaj et al. 2000). Thus, assessment of novelty-seeking (neophilic) behavior can reveal latent vulnerabilities to compulsive drug seeking (Carey et al. 2003; Laviola and Adriani 1998; Wingo et al. 2016). These preclinical findings appear to be borne out in human studies, in which people with substance use disorders tend to score higher on indices of neophilia than healthy subjects (Bardo et al. 1996; Bevins 2001; Wingo et al. 2016).

The canonical neuroanatomical substrate governing the behavioral effects of psychostimulants is the mesolimbic DA system (Baker et al. 2002; Di Chiara and Imperato 1988), which consists of midbrain DAergic neurons in the ventral tegmental area (VTA) and their target cells in the nucleus accumbens (NAcc) (Koob and Simon 2009; Nestler 2001). The medium spiny neurons (MSNs) of the NAcc also receive glutamatergic input from prefrontal cortex (PFC), hippocampus, thalamus, and amygdala (Baker et al. 2002; Koob and Volkow 2016). Within MSNs, simultaneous activation of D1Rs and NMDARs initiates a second messenger signaling cascade that converges on the Ras/Raf/MEK/ERK pathway (Baker et al. 2002; Berke and Hyman 2000; Lu et al. 2006), leading to activation of transcription factors which induce expression of immediate early genes (IEGs) (Lu et al. 2006; Sun et al. 2016). Indeed, MEK inhibitors block cocaine-induced locomotion (CIL) (Valjent et al. 2000), phosphorylation of ERK (pERK) (Valjent et al. 2006; Valjent et al. 2004), and expression of the IEG *c-fos* in the NAcc and amygdala (Valjent et al. 2000; Valjent et al. 2006; Valjent et al. 2004).

Behavioral and cellular effects of psychostimulants are modulated in part by regulators of G-protein signaling (RGS) proteins (Hooks et al. 2008; Sakloth et al. 2020). RGS proteins are characterized by an RGS domain that facilitates the termination of G-protein coupled receptor (GPCR) signaling by enhancing the GTPase activity of Ga subunits, limiting the duration of downstream signaling events (Ross and Wilkie 2000). This regulatory function of RGS proteins is particularly critical in the context of psychostimulant exposure, which induces excessive DA transmission and prolonged activation of DA-receptive GPCRs expressed by MSNs in the NAcc (Di Chiara and Imperato 1988; Gainetdinov et al. 2004).

RGS14 is a member of the R12 family of RGS proteins, sharing a conserved RGS domain that specifically catalyzes the GTPase activity of Gai/o family members (Hollinger et al. 2002; Sjögren and Neubig 2010). RGS14 also contains multiple unique functional domains that further diversify its range of intracellular signaling binding partners, enabling RGS14 to act as a scaffold for multiple signaling molecules such as certain active and inactive Ga subunits and small monomeric GTPases of the Ras/Rap subfamily family (Brown et al. 2015; Shu et al. 2007; Shu et al. 2010; Vellano et al. 2013). Importantly, RGS14 interactions with activated Ras suppress ERK signaling (Li et al. 2016; Shu et al. 2010; Willard et al. 2009). Although RGS14 is expressed in limbic regions (i.e., hippocampus, amygdala, and striatum) known to modulate responses to novelty and psychostimulants (Evans et al. 2014; Squires et al. 2020; Squires et al. 2018; Wingo et al. 2016), it has been studied almost exclusively in the context of hippocampal-dependent memory and synaptic plasticity (Lee et al. 2010; Squires et al. 2020), where it suppresses longterm potentiation (LTP) in CA2 pyramidal cells by regulating postsynaptic glutamatergic signaling, subsequent Ca^2+^ influx, and activation of CaMKII and ERK (Evans et al. 2018b; Lee et al. 2010).

Because of its expression pattern in the brain and capacity to suppress Ca^2+^-dependent ERK signaling, we hypothesized that RGS14 may regulate unconditioned locomotor responses and neuronal activity induced by cocaine and novelty. Thus, we assessed the consequences of genetic knockout of *Rgs14* on NIL, as well as CIL conducted in both familiar and novel environments. We also evaluated whether the absence of RGS14 protein affects neuronal activity, as measured by induction of c-fos or pERK (Lu et al. 2006), in brain regions that natively express RGS14 following exposure to cocaine in a novel environment (NIL + cocaine).

## Methods

### Subjects

*Rgs14* -/- (RGS14 KO) mice and their *Rgs14* +/+ (WT) littermate controls were maintained on a C57/BL6J background and genotyped by PCR, as previously described (Evans et al. 2018b; Lee et al. 2010). Behavioral experiments were conducted with adult (3-8 months old) mice of both sexes. All animal procedures and protocols were designed and performed in accordance with the National Institutes of Health Guidelines for the Care and Use of Laboratory Animals and were approved by the Emory University Institutional Animal Care and Use Committee. Mice were maintained on a 12 h light/12 h dark cycle with *ad libitum* access to food and water. All behavioral testing was conducted during the light cycle and analyses were conducted with experimenters blinded to genotype.

### Drugs

Cocaine hydrochloride (NIDA Drug Supply) was dissolved in sterile saline (0.9% NaCl) and injected intraperitoneally (i.p.) at a volume of 10 ml/kg. Sterile saline was injected to control for any effect of injection stress on behavior, and saline-treated animals were used as a control group for statistical comparison with cocaine-treated animals.

### Novelty-induced locomotion (NIL)

For all locomotor experiments, mice were placed into individual polycarbonate chambers (10” × 18” × 10”) surrounded by a 4×8 photobeam grid that recorded horizontal infrared beam breaks as a measure of ambulatory activity (Photobeam Activity System, San Diego Instruments, San Diego, CA). Two consecutive beam breaks were recorded as an ambulation, and total ambulatory activity was automatically sub-divided by location into two zones: the center or periphery of the chamber. A beam break was designated as “peripheral” if it was detected within 1 photobeam width (1.97”) from the walls of the test cage; all other ambulations were designated as “central.” For NIL, mice had never previously been exposed to the locomotor chambers, rendering the test environment novel. Novelty-induced ambulations were recorded for 1 h in 5-min bins, as previously described (Lustberg et al. 2020). Testing occurred in a brightly lit room where the animals were housed. Test cages were covered with a lid and contained a thin layer of standard bedding substrate.

### Cocaine-induced locomotion (CIL)

RGS14 KO mice and WT littermates were used to generate a within-subjects dose response curve of acute, cocaine-induced locomotion. Mice received a single dose per test session, and test days were staggered by 1 week to prevent sensitization, as described (Manvich et al. 2019; Weinshenker et al. 2002). On each test day, mice were habituated to the locomotor chambers for 30 min, during which baseline ambulations were recorded. Mice were then briefly removed from their chambers and given an injection of either saline or cocaine (5, 10, or 20 mg/kg). Mice were returned to their chambers, and locomotor activity was recorded for an additional 1 h in 5 min bins. All mice received all doses over the duration of the experiment. Dose order was pseudorandomized to avoid order effects.

### Cocaine-induced locomotion in a novel environment (NIL + cocaine)

A cohort of mice used for the CIL dose-response experiment was allowed a 4-week washout period prior to NIL with acute cocaine pre-treatment (NIL + cocaine). From pilot experiments and previous studies (Manvich et al. 2019; Porter-Stransky et al. 2019), we determined that mice retain memory for a novel environment for less than 1 week. All mice were injected with 20 mg/kg cocaine and immediately placed into a locomotor chamber for 1 h, during which time ambulations were recorded in 5 min bins as before. Mice were left undisturbed in their chambers until 90 min had elapsed, after which mice were euthanized for c-fos and pERK immunohistochemistry.

### Immunohistochemistry (IHC)

Following the NIL + cocaine test (90 min after test onset), mice were euthanized with an overdose of sodium pentobarbital (Fatal Plus, 150 mg/kg, i.p.; Med-Vet International, Mettawa, IL), and transcardially perfused with cold 4% paraformaldehyde in 0.01 M PBS. After extraction, brains were post-fixed for 24 h in 4% paraformaldehyde at 4°C, and then transferred to 30% sucrose/PBS solution for 72 h at 4°C. Brains were embedded in OCT medium (Tissue-Tek; Sakura, Torrance, CA) and serially sectioned by cryostat (Leica) into 40-μm coronal slices spanning the striatum through the hippocampus. Brain sections were stored in 0.01 M PBS (0.02% sodium azide) at 4°C before IHC.

For c-fos IHC, brain sections were blocked for 1 h at room temperature in 5% normal goat serum (NGS; Vector Laboratories, Burlingame, CA) diluted in 0.01 M PBS/0.1% Triton-X permeabilization buffer. Sections were then incubated for 48 h at 4°C in NGS blocking/permeabilization buffer, including a primary antibody raised against c-fos (rabbit anti-c-fos, Millipore, Danvers, MA, ABE457; 1:5000). After washing in 0.01 M PBS, sections were incubated for 2 h in blocking/permeabilization buffer with goat anti-rabbit AlexaFluor 488 (Invitrogen, Carlsbad, CA; 1:500).

For pERK and RGS14 IHC, brain sections were first subjected to antigen retrieval in 10 mM sodium citrate buffer (3 min at 100°C) prior to blocking, as described above. Sections were then incubated for 24 h at 4°C in NGS blocking/permeabilization buffer, including primary antibodies raised against pERK (rabbit anti-Phospho-p44/42 MAPK, Cell Signaling Technology, Danvers, MA, #4370; 1:1000) or RGS14 (mouse anti-RGS14, NeuroMab, Davis, CA, clone N133/21; 1:500). The sections were then incubated for 2 h in blocking/permeabilization buffer including goat anti-rabbit AlexaFluor 568 or goat anti-mouse 488 (Invitrogen; 1:500). Sections were washed in 0.01 M PBS after primary and secondary antibody incubations, and then mounted onto Superfrost Plus slides (Thermo Fisher Scientific, Waltham, MA). Once dry, slides were coverslipped with Fluoromount-G + DAPI (Southern Biotech, Birmingham, AL).

### Fluorescent imaging and quantification

Fluorescent micrographs of immunostained sections were acquired on a Leica DM6000B epifluorescent upright microscope at 10x or 20x magnification for regional RGS14 characterization, and at 10x magnification with uniform exposure parameters for regional c-fos and pERK quantification. For c-fos and pERK quantification, atlas-matched sections were selected from each animal at the level of the NAcc and dorsal hippocampus. A standardized region of interest was drawn for all images to delineate the borders of discrete structures in all subjects. The structures selected for comparison were the NAcc (shell and core), central amygdala (CeA), piriform cortex (Pir Ctx), and dorsal hippocampal subfields CA1, CA2, and CA3.

Image processing and analysis were performed using ImageJ software. The analysis pipeline included background subtraction, intensity thresholding (Otsu method), and automated cell counting within defined regions of interest, guided by automated size and shape criteria for c-fos+ and pERK+ cells (size: 70–100 pixel^2^, circularity: 0.6–1.0).

### Statistical analysis

For locomotor experiments, within-session ambulations and dose-response curves were compared by two-way repeated measures ANOVA with Sidak’s post hoc test. Total ambulations between genotypes were evaluated with unpaired *t*-tests. Central and peripheral ambulations were compared between genotypes using a two-way ANOVA with Sidak’s post hoc test. For the NIL and NIL + cocaine experiment, thigmotaxis ratios (center ambulations / peripheral ambulations) were compared between genotypes across experimental conditions using a twoway ANOVA with Tukey’s post hoc test.

For c-fos and pERK quantification, genotype differences were compared in the seven regions mentioned above following the NIL + cocaine test. Comparisons were made within regions and between genotypes by *t*-tests using the Holm-Sidak correction for multiple comparisons. The threshold for adjusted significance was set at *p* < 0.05, and two-tailed variants of tests were used throughout. The statistical analyses were conducted and graphs were designed using Prism v8 (GraphPad Software, San Diego, CA).

## Results

### RGS14 is expressed in discrete cortical and limbic structures sensitive to novelty and cocaine

Immunohistochemical detection of RGS14 immunoreactivity (RGS-ir) in coronal brain sections from RGS14 WT mice revealed RGS14-ir in dorsal hippocampal subfields CA1 and CA2, CeA, Pir Ctx, NAcc core, and NAcc shell (Fig. 1a, b). RGS14-ir was absent in dorsal hippocampal subfield CA3 (Fig. 1c). CA3 pyramidal cells are activated by psychostimulants (Luo et al. 2011) and are involved in novelty detection (He et al. 2002; Wagatsuma et al. 2018); thus, we selected CA3 as a RGS14-negative control region for comparison with RGS14-ir regions. In CA1, Pir Ctx, and NAcc, RGS14-ir was detected predominantly in neurites as previously described (Squires et al. 2018) while intense RGS14-ir was observed in both cell bodies and neurites of CA2 (Carstens and Dudek 2019; Lee et al. 2010) and CeA neurons (Fig. 1b,c).

**Figure 1.**
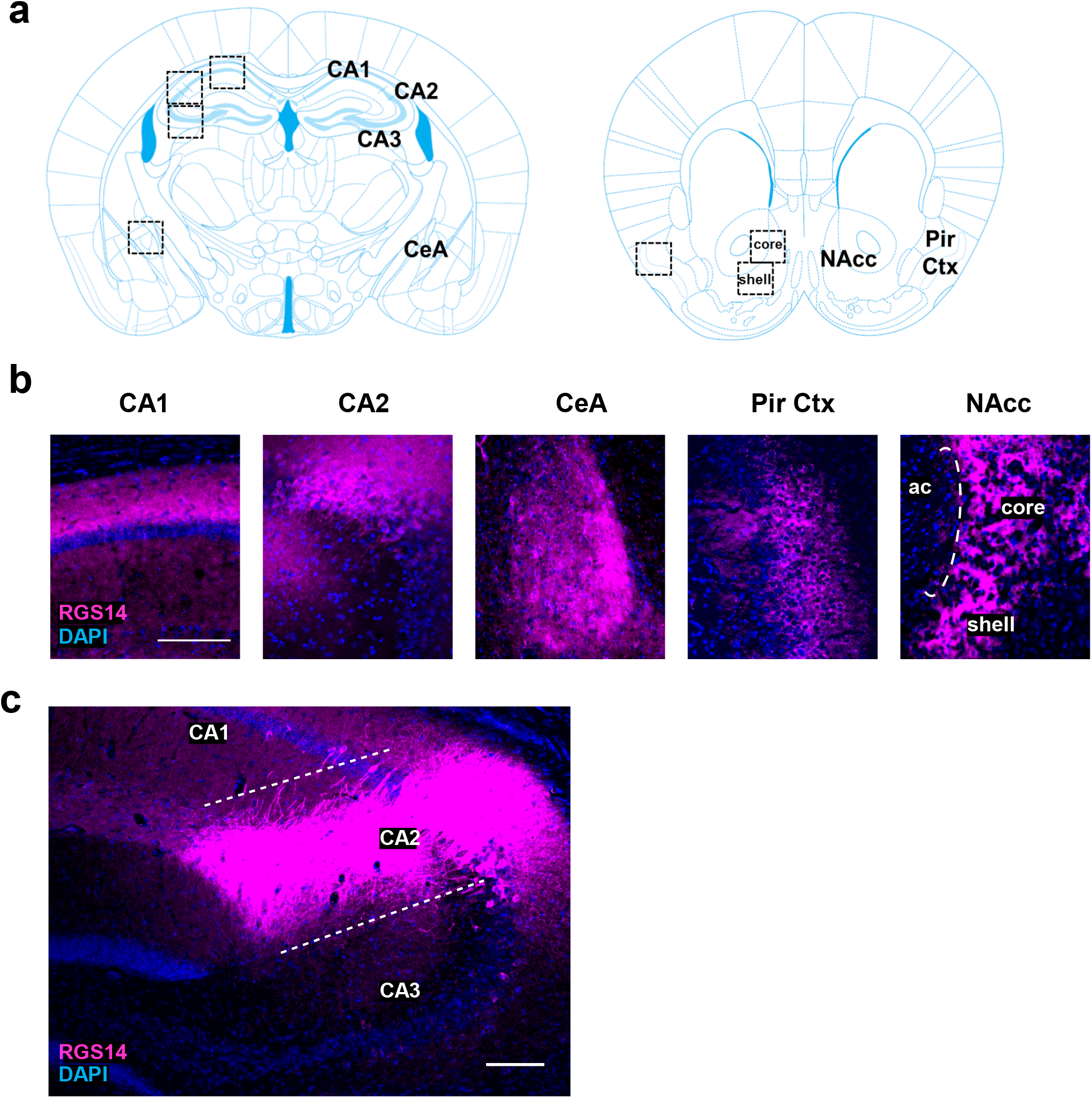
Detection of RGS14 immunoreactivity (RGS-ir) in cortical and limbic structures of the mouse brain. **a** Brain regions selected for immunohistochemical analysis of endogenous RGS14-ir, as well as c-fos and pERK induction following exposure to novelty and cocaine. **b** RGS14 (magenta) is endogenously expressed in the dorsal hippocampus areas CA1 and CA2, central amygdala (CeA), piriform cortex (Pir Ctx), and nucleus accumbens (NAcc) core and shell subregions. The anterior commissure (ac) is depicted to delimit the NAcc. **c** There is no endogenous RGS14-ir within the CA3 subfield of the hippocampus of WT mice. Scale bars denote 100 μm.

Though the behavioral functions of CA2 have been studied mostly in the context of social memory (Dudek et al. 2016; Hitti and Siegelbaum 2014; Meira et al. 2018), the remaining RGS14-ir regions have established roles in spatial memory (McDonald and White 2013), novelty detection (Moreno-Castilla et al. 2017; Strauch and Manahan-Vaughan 2020), motivation (Baker et al. 2002), and behavioral responses to stress or psychostimulants (Fadok et al. 2018; Kabbaj et al. 2000). The expression pattern of RGS14 throughout these distributed brain regions, all of which receive glutamatergic and DAergic innervation (Datiche and Cattarelli 1996; Fadok et al. 2018; McNamara and Dupret 2017; Takeuchi et al. 2016), guided our decision to assess unconditioned behavioral responses to novelty and psychostimulants in RGS14 KO mice, which to date have been almost exclusively examined in behavioral paradigms that assess learning and memory.

### RGS14 deficiency attenuates novelty-induced locomotion (NIL) but increases thigmotaxis

To determine the effect of RGS14 deficiency on innate novelty-induced exploratory behavior, a cohort of age- and sex-matched WT and RGS14 KO mice were compared in the NIL test. Total ambulations, center ambulations, and peripheral ambulations were measured in 5-min bins over 1 h. WT and RGS14 KO mice displayed similar initial novelty detection, with the greatest amount of locomotor activity occurring at the beginning of the task (Fig. 2a). A two-way repeated measures ANOVA (genotype x time) showed significant main effects for genotype (*F*(1, 37) = 7.07, *p* = 0.012) and time (*F*(4.733, 175.1) = 55.72, p < 0.0001) with no significant interaction (*F*(11, 407) = 1.036, *p* > 0.05) (Fig. 2a). The genotype differences in locomotion emerged in the second half of the NIL test but did not reach statistical significance (*p* > 0.05) at any single time point when analyzed with Sidak’s multiple comparison tests.

**Figure 2.**
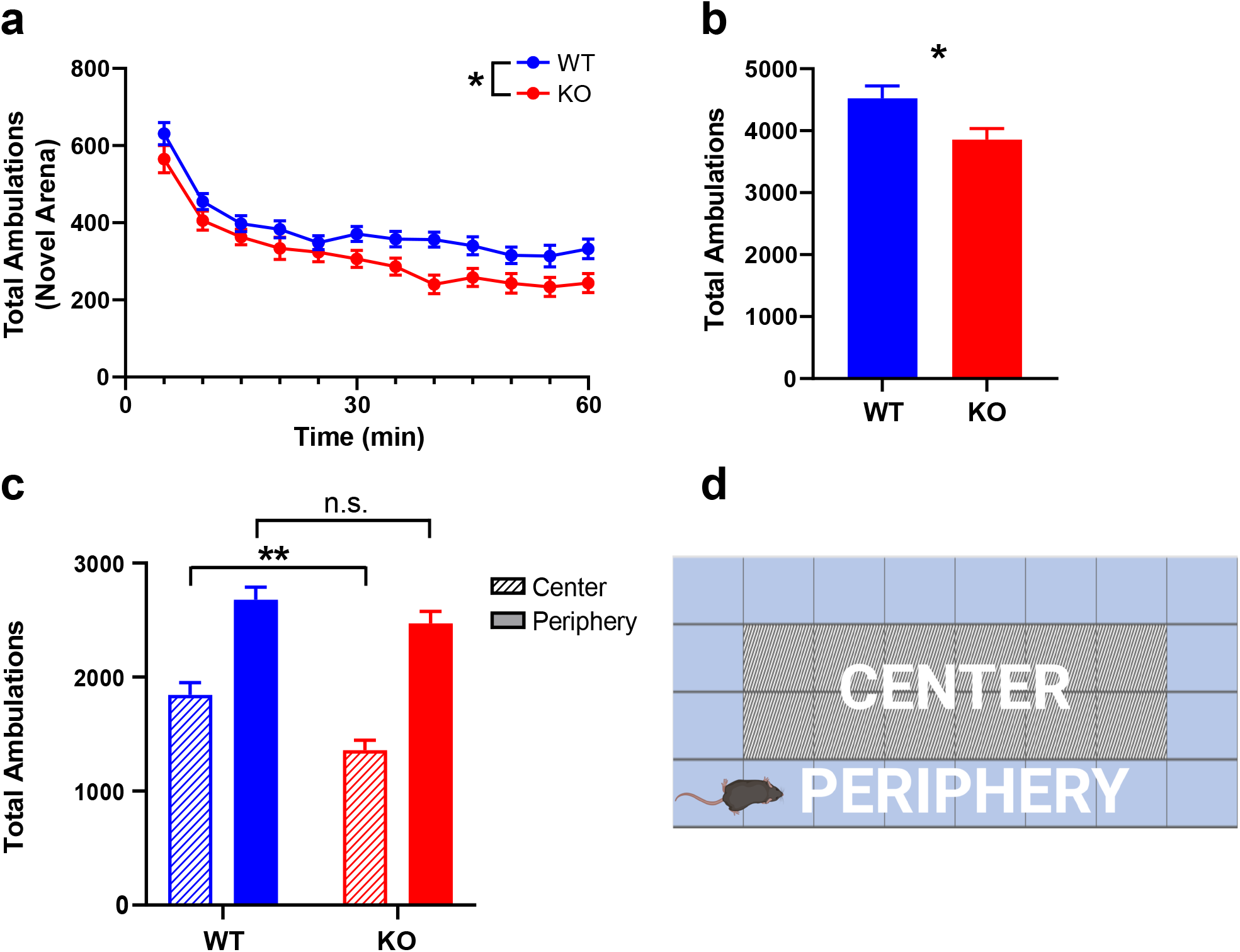
RGS14 KO mice exhibit attenuated novelty-induced locomotion (NIL) and increased thigmotaxis but show no deficits in novelty detection or within-trial habituation. **a** Locomotor activity measured in 5-min bins over 1 h in a novel environment for WT (blue) and KO (red) mice. Ambulations decreased over time for both genotypes as mice habituated to the novel environment, but the reduction in activity was greater in KO compared to WT. **b** KO mice exhibited fewer total ambulations than WT during the NIL test. **c** While peripheral ambulations were similar between WT (blue, solid bar) and KO mice (red, solid bar), KO mice made fewer center ambulations (red, striped bar) than WT (blue, striped bar), indicative of thigmotaxis. **d** Schematic of locomotor testing chamber with delineated central and peripheral regions. n = 19 – 20 per genotype. n.s. = not significant, \**p* < 0.05, \*\**p* < 0.01. *Created with BioRender.com*.

An unpaired t-test of total ambulations in the NIL paradigm revealed that RGS14 KO mice were hypoactive in novel environments compared to WT controls (3858 ± 785 vs 4524 ± 875; *t*(37) = 2.50, *p* = 0.02) (Fig. 2b). Because the hypoactivity exhibited by RGS14 KO mice appeared to be driven by thigmotaxis behavior (reduced locomotion in the center of the environment), a twoway ANOVA (genotype x area) of peripheral and center ambulations was performed, which revealed a main effect of genotype (*F*(1, 74) = 11.46, *p* = 0.0011) and area (*F*(1, 74) = 90.28, *p* < 0.0001) (Fig. 2c). Tukey’s post hoc test showed that while peripheral ambulations did not differ by genotype (*p* > 0.05) center ambulations were significantly lower in RGS14 KO mice compared to WT controls (*p* < 0.01). Collectively, these findings indicate that RGS14 KO mice exhibit reduced locomotion in a novel environment, characterized by increased thigmotaxis.

### RGS14 deficiency augments cocaine-induced locomotion (CIL) in a familiar environment

Given that RGS proteins modulate psychostimulant-induced locomotion (Rahman et al. 2003), and that RGS14 suppresses ERK-signaling which is critical for cocaine-mediated plasticity and cellular activation (Lu et al. 2006; Shu et al. 2010), we sought to determine the impact of RGS14 deletion on CIL. To assess CIL in a familiar environment, mice were habituated to the testing chamber for 30 min prior to saline or cocaine administration. An unpaired t-test of total ambulations following saline injection was performed and indicated no baseline difference between genotypes that could confound interpretation of the CIL test, (*t*(26) = 0.63, *p* > 0.05) (Fig. 3a). Additionally, a two-way ANOVA (genotype x area) of central and peripheral ambulations after saline injection showed a main effect of area (*F*(1, 52) = 12.92, *p* = 0.0007) but not genotype (*F*(1, 52) = 0.73, *p* > 0.05) or interaction (*F*(1, 52) = 0.01, *p* > 0.05), indicating that there are no baseline differences in thigmotaxis between strains when assessed in a familiar environment without psychostimulants (Fig. 3b). A two-way repeated measures ANOVA of cocaine dose-response curves indicated a main effect of dose (*F*(3,78) = 90.99, *p* < 0.0001) but not genotype (*F*(1,26) = 2.841, *p* > 0.05), with a dose x genotype interaction (*F*(3,78) = 3.60, *p* = 0.02) (Fig. 3c). However, Sidak’s post hoc test did not indicate significant differences between genotypes at any single dose.

**Figure 3.**
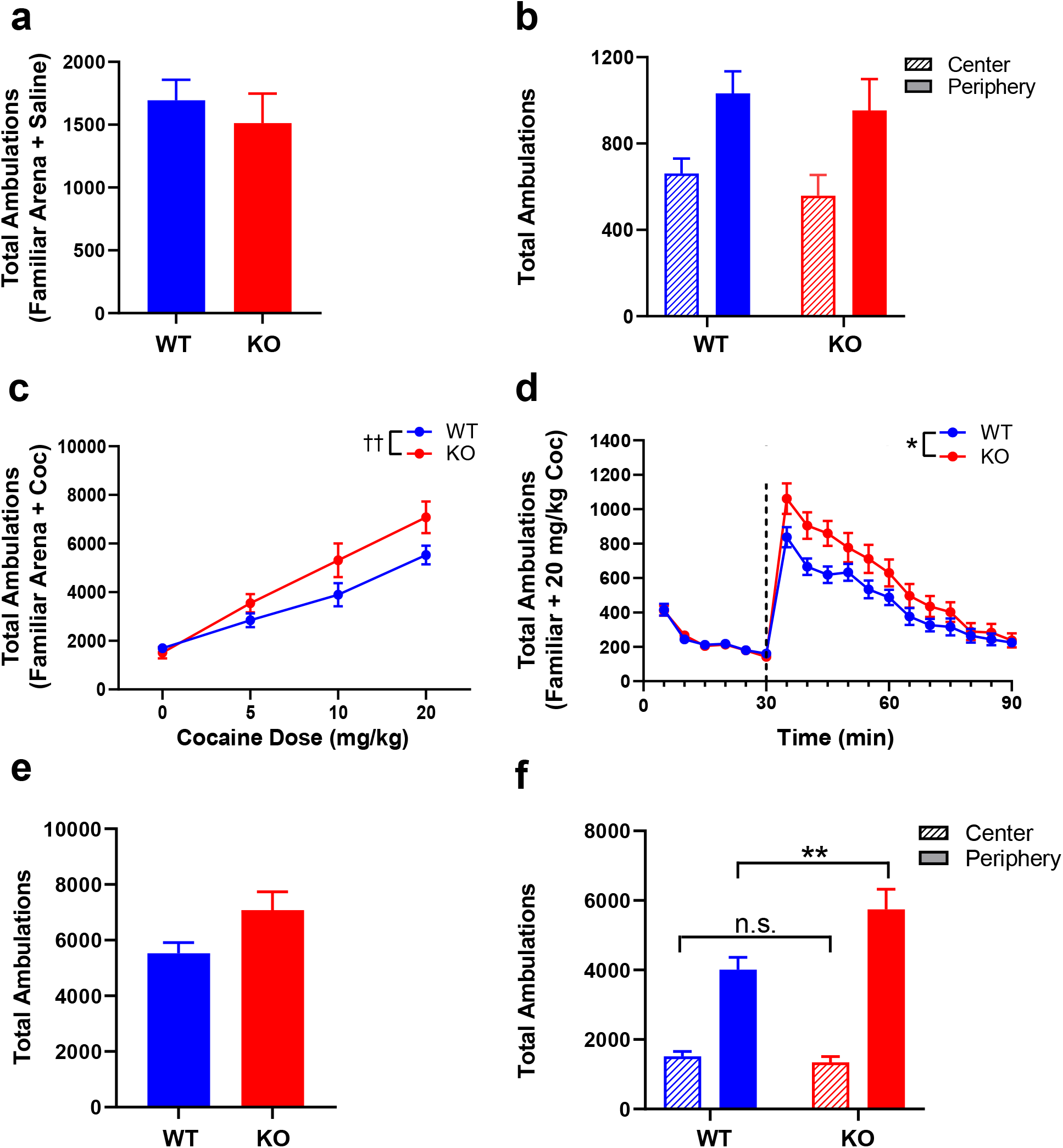
RGS14 KO mice exhibit enhanced cocaine-induced locomotion (CIL) and thigmotaxis in a familiar environment. **a** WT (blue) and KO (red) mice exhibited similar total locomotor activity during 1 h in a familiar environment following saline injection. **b** Both genotypes exhibited comparable levels of peripheral (solid bars) and central (striped bars) ambulations in a familiar environment. **c** Dose-response curves of acute cocaine-induced locomotion for WT (blue) and KO (red) in a familiar environment showed a significant dose x genotype interaction, evident at higher doses of cocaine. **d** Locomotor activity during a 30-min habituation period and 1 h following administration of 20 mg/kg cocaine (dashed line). KO mice displayed an enhanced locomotor response to cocaine treatment. **e** Total ambulations after 20 mg/kg cocaine were increased among KO mice compared to WT but did not reach significance. **f** Cocaine in a familiar environment had no effect on center ambulations (WT = blue, striped bar; KO = red, striped bar) but did increase peripheral ambulations in KO mice (red, solid bar) compared to WT (blue, solid bar). n = 14 per genotype. n.s. = not significant, \**p* < 0.05, ** *p* < 0.01, ^††^ *p* < 0.05 for the effect of interaction.

Because genotype differences appeared most pronounced at the 20 mg/kg cocaine dose, we then evaluated within-session locomotor activity at this dose. A two-way repeated measures ANOVA of ambulations within the first 30 min following cocaine administration showed main effects of genotype (*F*(1, 26) = 5.159, *p* = 0.032) and time (*F*(2.293, 59.62) = 34.16, p < 0.0001), but no genotype x time interaction (*F*(5, 130) = 0.9633, *p* > 0.05) (Fig. 3d). Sidak’s post hoc tests did not indicate significant differences between genotypes at any individual time point. While total ambulations at the 20 mg/kg dose only showed a trend for significance via an unpaired t-test (*t*(26) = 2.049, *p* = 0.051) (Fig. 3e), a two-way ANOVA (genotype x area) of central and peripheral ambulations at this dose showed main effects of genotype (*F*(1, 52) = 4.684, *p* = 0.04) and area (*F*(1, 52) = 91.73, *p* < 0.001), as well as a genotype x area interaction (*F*(1, 52) = 7.063, *p* = 0.01) (Fig. 3f). Sidak’s post hoc test indicated that although RGS14 KO and WT mice exhibited similar central ambulations (*p* > 0.05), peripheral ambulations were significantly higher in the RGS14 KO mice (*p* < 0.01). These results indicate that cocaine-induced locomotion in a familiar environment is augmented in RGS14 KO mice, specifically by promoting thigmotaxis (enhancing locomotion in the periphery).

### Cocaine augments novelty-induced locomotion in RGS14-deficient mice but increases thigmotaxis

Given the surprising finding that RGS14 KO mice show opposite locomotor responses to novelty and cocaine, which are usually positively correlated, we assessed the effects of cocaine treatment on NIL. Previous studies have demonstrated that high doses of cocaine (≥ 10 mg/kg) increase locomotor activity in novel environments but also promote anxiety-like behavior, such as thigmotaxis (Blanchard and Blanchard 1999; Schank et al. 2008; Simon et al. 1994). We reasoned that genotype differences in thigmotaxis behavior would be maximally apparent when high dose cocaine was paired with novelty exposure (Blanchard and Blanchard 1999; Simon et al. 1994). Thus, we performed a test in which 20 mg/kg cocaine was administered immediately before placing mice in the novel test cage environment.

Total ambulations, center ambulations, and peripheral ambulations were measured in 5-min bins across 1 h. A two-way repeated measures ANOVA (genotype × time) showed a main effect of genotype (*F*(1,12) = 7.70, *p* = 0.02) and time (*F*(3.22, 38.63) = 29.49, *p* <0.0001) (Fig. 4a). Although there was no overall time x genotype interaction (*F*(11, 132) = 1.34, *p* > 0.05), Sidak’s post hoc tests revealed that significant genotypes differences in locomotion emerged early in the NIL + cocaine test rather than late, with RGS14 KO mice exhibiting more activity than WT at the 15-min (*p* = 0.02) and 25-min (*p* = 0.04) time points (Fig. 4a).

**Figure 4.**
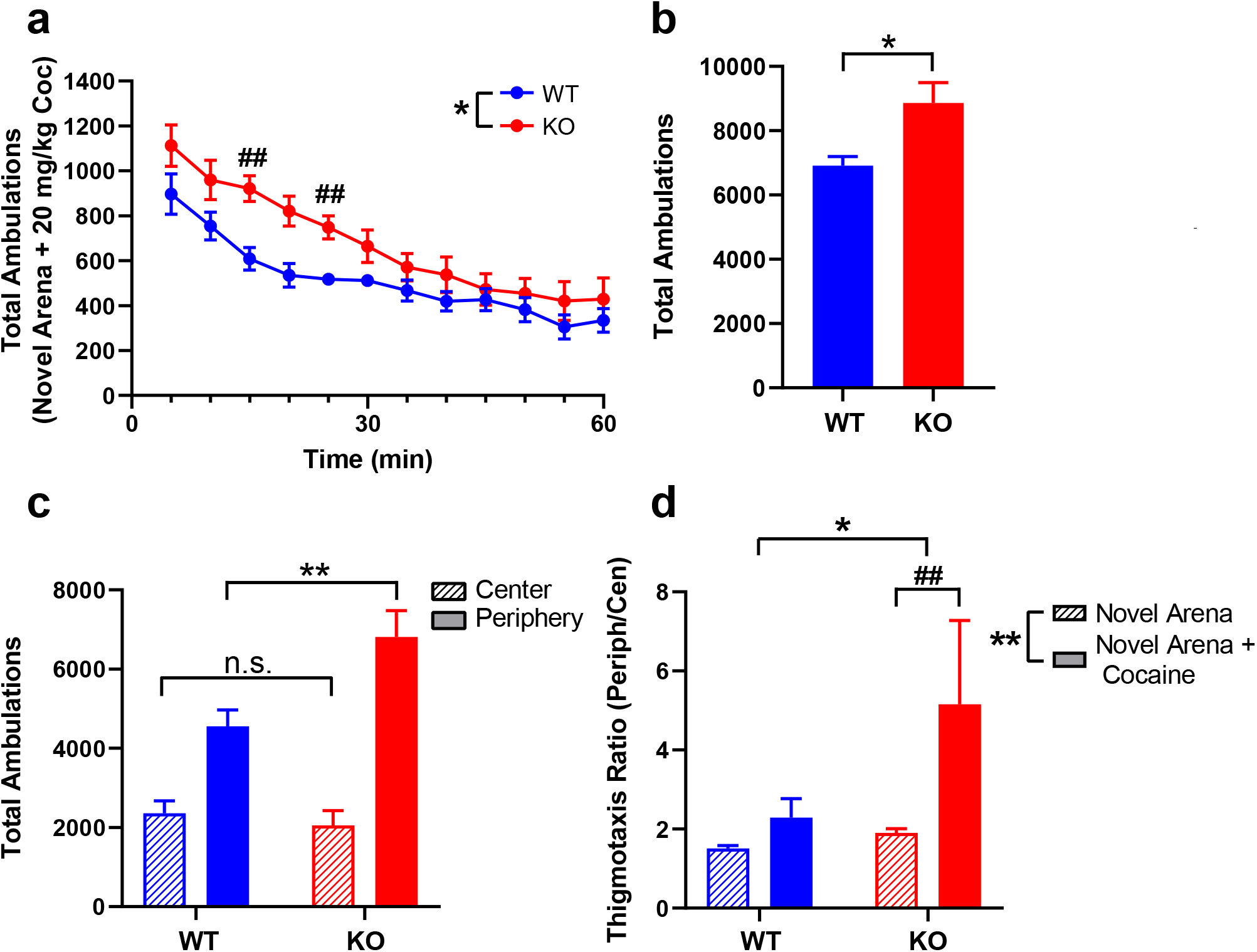
RGS14 KO mice are hypersensitive to cocaine-induced locomotion and thigmotaxis in a novel environment. **a** Locomotor activity in WT (blue) and KO mice (red) over 1 h following administration of 20 mg/kg cocaine in a novel environment. Ambulations were significantly different between genotypes, with KO mice exhibiting higher activity than WT at 15 and 25 min following cocaine administration. **b** Cocaine increased total ambulations in RGS14 KO mice (red) significantly more than WT (blue). **c** Cocaine did not affect central ambulations between genotypes (WT = blue, striped bar; KO = red, striped bar) but increased peripheral ambulations in RGS14 KO mice (red, solid bar) compared to WT (blue, solid bar). **d** Compared to the novel environment alone, cocaine in a novel environment selectively increased thigmotaxis ratio in RGS14 KO (red), but not WT (blue) mice. n = 7 per genotype. n.s. = not significant, \**p* < 0.05, ** *p* < 0.01, ## p < 0.05 by post hoc comparison.

An unpaired t-test of total ambulations in the NIL + cocaine test revealed that RGS14 KO mice were hyperactive, rather than hypoactive, in novel environments relative to WT controls (8866 ±1620 vs 6910 ± 747; *t*(12) = 2.83, *p* =0.02) (compare Fig. 4b with Fig. 2b). A two-way ANOVA (genotype x area) of central and peripheral ambulations showed main effects of genotype (*F*(1, 24) = 4.461, *p* < 0.05) and area (*F*(1, 24) = 56.36, *p* < 0.001), as well as a genotype x area interaction (*F*(1,24) = 7.649, *p* = 0.01) (Fig. 4c). Sidak’s post hoc test revealed similar measures of center ambulations between genotypes (*p* > 0.05), but significantly increased peripheral ambulations in RGS14 KO mice (*p* < 0.001).

To compare thigmotaxis behavior across tests, we computed a thigmotaxis ratio (peripheral ambulations / center ambulations) for each genotype in the NIL and NIL + cocaine tests. A two-way ANOVA (genotype x condition) revealed a main effect of genotype (*F*(1, 49) = 6.57, *p* = 0.01) and condition (*F*(1, 49)= 10.04, *p* <0.01), and a strong trend for a genotype x condition interaction (*F*(1, 49) = 3.70, *p* = 0.06) (Fig. 4d). Tukey’s post hoc tests revealed that the thigmotaxis ratio for RGS14 KO mice was significantly higher in the NIL + cocaine test compared to NIL alone (*p* < 0.01), while thigmotaxis ratios did not differ between NIL and NIL + cocaine in WT mice (*p* > 0.05).

### RGS14 deficiency enhances induction of c-fos and pERK in the hippocampus and amygdala

Phosphorylation of ERK (pERK) in response to synaptic signaling regulates postsynaptic plasticity and promotes the upregulation of immediate early genes (IEG) such as c-fos (Lu et al. 2006), and RGS14 has previously been shown to block ERK signaling (Li et al. 2016; Shu et al. 2010) To identify the impact of RGS14 on neuronal activity and signaling induced by cocaine + novelty exposure, RGS14 KO mice and WT littermates were euthanized 90 min after the onset of NIL + cocaine test for quantification of c-fos+ and pERK+ cells in six brains regions that express RGS14 under normal conditions and one negative control region that does not express RGS14 (Fig. 1).

Following combined exposure to novelty and cocaine, RGS14 KO and WT mice demonstrated similar levels of c-fos in CA3 (*t*(12) = 0.75, *p* > 0.05), Pir Ctx (*t*(12) = 0.59, *p* > 0.05), the NAcc core (*t*(12) = 0.19, *p* > 0.05), and the NAcc shell (*t*(12) = 0.56, *p* > 0.05) (Fig. 5a). However, RGS14 KO mice showed increased c-fos induction in CA1 (*t*(12) = 4.83, *p* < 0.01), CA2 (*t*(12) = 5.49, *p* < 0.01), and the CeA (*t*(12) = 7.60, *p* < 0.001) compared to WT, suggesting enhanced neuronal activation in these regions in the absence of RGS14 (Fig. 5a).

**Figure 5.**
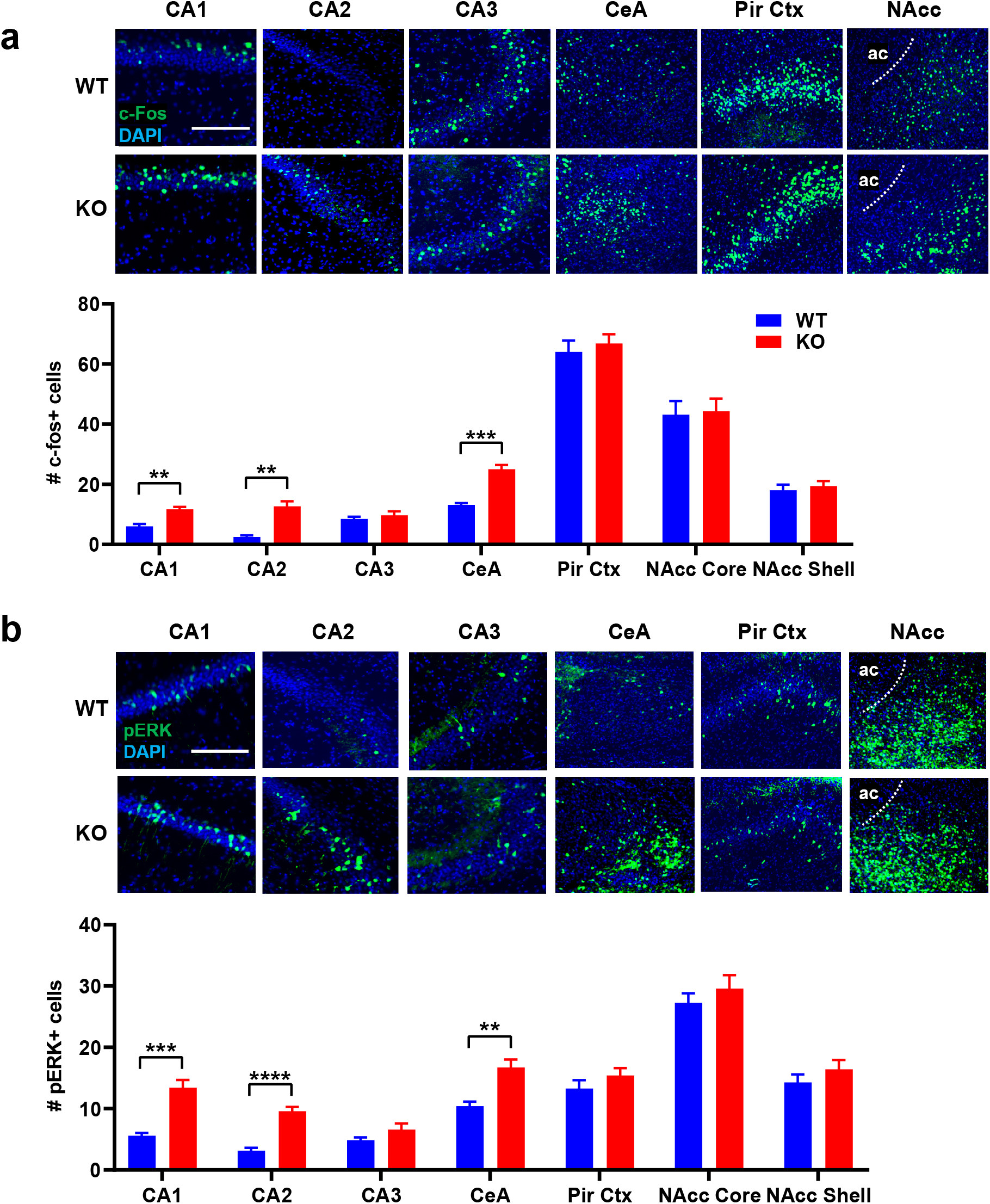
RGS14 deficiency enhances expression of c-fos and pERK in select brain regions following cocaine administration and exposure to a novel environment. **a** c-fos induction was increased in CA1, CA2, and CeA of RGS14 KO mice (n = 7; bottom row) compared to WT mice (n = 7; top row) following NIL + cocaine, but there were no genotype differences in c-fos induction in CA3, Pir Ctx, or NAcc. **b** pERK expression was increased in CA1, CA2, and CeA of KO mice (n = 7; bottom row) compared to WT mice (n = 7; top row) following NIL + cocaine, but there were no genotype differences in pERK expression in CA3, Pir Ctx, or NAcc. Scale bars denote 100 μm. ** *p* < 0.01, *\*\*\*p* < 0.001, *\*\*\*\*p* < 0.0001.

An identical pattern was observed for regional analysis of pERK+ immunoreactive cells. Following exposure to novelty + cocaine, RGS14 KO and WT mice displayed comparable levels of pERK in CA3 (*t*(12) = 1.5, *p* > 0.05), the Pir Ctx (*t*(12) = 1.18, *p* > 0.05), the NAcc core (*t*(12) = 0.85, *p* > 0.05), and the NAcc shell (*t*(12) = 1.07, *p* > 0.05) (Fig. 5b), while pERK immunoreactivity was augmented in RGS14 KO mice within CA1 (*t*(12) = 5.79, *p* < 0.001), CA2 (*t*(12) = 7.54, *p* < 0.0001), and the CeA (*t*(12) = 4.22, *p* < 0.01) (Fig. 5b).

## Discussion

### RGS14 is expressed in limbic subregions activated by novelty and psychostimulants

In this study, we observed RGS14-ir in neurons within hippocampal area CA1 and CA2, Pir Ctx, NAcc core and shell, and CeA. RGS14 expression in the brain is highest in CA2 pyramidal cells (Evans et al. 2014; Gerber et al. 2019), where it suppresses synaptic plasticity (Evans et al. 2018b; Lee et al. 2010). Indeed, CA2 neurons display enhanced excitability and physiologically abnormal plasticity at CA3 → CA2 synapses in the absence of RGS14 (Evans et al. 2018b; Lee et al. 2010; Zhao et al. 2007), and RGS14 KO mice demonstrate superior spatial and object recognition memory relative to WT mice (Lee et al. 2010). Interestingly, overexpression of RGS14 in CA1 neurons blocks synaptic plasticity at CA3 → CA1 synapses (Evans et al. 2018b), which is readily induced under normal conditions (Carstens and Dudek 2019), and suggests that high levels of RGS14 suppress plasticity in CA2 through signaling mechanisms shared by CA1 (Evans et al. 2018; Lee et al. 2010).

The robust RGS14-ir we saw in this study within CeA represents the first report of RGS14 expression in the amygdala of rodents, and may provide insight into previous behavioral studies of RGS14 KO mice. Recently, female RGS14 KO mice were reported to exhibit enhanced cued fear memory relative to WT controls (Alexander et al. 2019). The authors of this study attributed this sex-specific facilitation of fear learning to the absence of RGS14 in CA2. While this is one possibility, an alternative explanation is that this phenotype involves the absence of RGS14 in CeA, rather than the dorsal hippocampus. In fact, cued fear learning does not require the dorsal hippocampus and depends instead on the integrity of the amygdala (McDonald and White 2013; Phillips and LeDoux 1992), including the CeA, which we show here endogenously expresses RGS14. Our results also highlight candidate brain regions for further exploration of RGS14 function. Neurons in CeA, Pir Ctx, and NAcc are capable of synaptic plasticity and are activated by novelty or psychostimulants (Fadok et al. 2018; Russo et al. 2010; Wilson et al. 2004), but the behavioral and physiological functions of RGS14 in these regions remain unknown.

### RGS14 KO mice exhibit reduced locomotion but enhanced thigmotaxis in response to novelty

Our findings indicate that while RGS14 KO mice do not differ from WT in novelty detection or habituation during the NIL test, they exhibit a reduction in overall locomotion driven by enhanced thigmotaxis. Usually, NIL behavior is interpreted as a measure of neophilia, with hypoactive responses indicating decreased exploratory drive (Bevins 2001; Wingo et al. 2016). On the other hand, exposure to a novel environment is stressful; NIL can also be interpreted as an assay of novelty-induced anxiety (neophobia), with low locomotor responses indicating greater neophobia (Griebel et al. 1993; Walker et al. 2009).

We argue that total ambulatory activity alone cannot discern neophilic versus neophobic responses, since anxiogenic and anxiolytic drugs have variable effects on gross NIL responses (Angrini et al. 1998; Matsubara and Matsushita 1982; Simon et al. 1994). Further, we posit that total locomotor activity in the NIL or open field test is not a reliable measure of anxiety (Fraser et al. 2010). Our findings and those of other groups (Matsubara and Matsushita 1982; Simon et al. 1994) suggest that gross locomotor activity in a novel environment has poor affective resolution (Crawley 2007); instead, the valence of behavioral responses to novel environments should be interpreted using additional metrics that account for biases toward thigmotaxis and changes in types of locomotor activity over time (Lustberg et al. 2020; Simon et al. 1994). These same standards should also be applied when measuring the locomotor effects of drugs in familiar environments.

Importantly, anxiolytic drugs consistently *increase* thigmotaxis in novel environments, while anxiolytic drugs consistently *decrease* thigmotaxis in novel environments (Fraser et al. 2010; Simon et al. 1994). To refine our characterization of NIL behavior, we also assessed genotype differences in thigmotaxis, which were marked. We encourage researchers examining locomotor phenotypes in novel environments to consider multiple variables when interpreting behavioral findings, including task duration, total ambulations, thigmotaxis ratio, initial novelty response, and within-trial habituation. These additional measures may help identify previously unappreciated locomotor phenotypes, explain unexpected behavioral differences, and foster a more nuanced interpretation of experimental findings.

While we cannot definitively conclude whether the NIL deficit in RGS14 KO mice reflects reduced neophilia or increased neophobia, the hypoactive NIL response combined with augmented thigmotaxis in RGS14 KO mice is more consistent with the latter interpretation. To our knowledge, the present study is the first to identify a behavioral impairment in RGS14 KO mice, which outperform WT controls in tests of spatial and emotional memory (Alexander et al. 2019; Lee et al. 2010).

### RGS14 KO mice demonstrate increased locomotion and thigmotaxis in response to cocaine

NIL and CIL responses are often positively correlated, with high NIL responders being more sensitive to rewarding and habit-forming psychostimulant effects (Bardo et al. 1996; Wingo et al. 2016). In contrast to the hypoactivity they exhibited in NIL, RGS14 KO mice displayed enhanced locomotion and thigmotaxis when administered high-dose cocaine (≥ 10 mg/kg) in a familiar environment. Though the phenotypes of RGS14 KO mice in NIL and CIL were inversely related (Carey et al. 2003; Walker et al. 2009), both novelty- and cocaine-induced thigmotaxis were enhanced in RGS14 KO mice relative to WT controls, suggesting that mice may be more sensitive to neophobia and cocaine-induced anxiety in the absence of RGS14 (Griebel et al. 1993; Simon et al. 1994).

Interestingly, the locomotor deficit in RGS14 KO mice during NIL emerged in the second half of the 1-h test, while the locomotor enhancement in CIL emerged in the first half of the test. Thus, the “delayed” hypoactivity in RGS14 KO mice in the NIL test may represent the effects of RGS14 disruption on signaling pathways that are distinct from those that mediate their “early” hyperactivity in CIL. Yet another possibility is that RGS14 binding interactions differ within each brain region where it is expressed, or that RGS14 signaling varies as a function of other factors like stimulus intensity or context familiarity.

### Cocaine enhances locomotor activity and thigmotaxis in RGS14 KO mice exposed to a novel environment

NIL and CIL testing are typically performed in unfamiliar and familiar environments, respectively, to distinguish the locomotor-activating effects of novelty and cocaine (Fraser et al. 2010; Walker et al. 2009). Given that the RGS14 KO mice displayed hypoactive NIL and hyperactive CIL but increased thigmotaxis in both assays, we reasoned that pretreating RGS14 KO mice with cocaine prior to exposing them to the novel test cage would further enhance thigmotaxis (Simon et al. 1994). Indeed, RGS14 KO mice were hyperactive in the NIL + cocaine test, demonstrating more locomotor activity than WT controls in the novel test cage during the first half of the 1 h testing session. As in the CIL and NIL tests, RGS14 KO mice also demonstrated more thigmotaxis than WT controls in the NIL + cocaine test. Interestingly, when we compared thigmotaxis ratios between genotypes and across the NIL or NIL + cocaine conditions, we discovered that cocaine selectively enhanced the thigmotaxis ratio in RGS14 KO mice.

These behavioral findings can be interpreted in at least three ways. The first explanation is that RGS14 KO mice are less neophilic, and that cocaine-induced increases in catecholamine and glutamate signaling enhance neophilia in RGS14 KO mice during NIL + cocaine testing beyond the level of WT controls. This explanation seems unlikely because RGS14 KO and WT mice did not differ in their initial novelty responses during the NIL test, suggesting that neophilia is unaffected by the absence of RGS14. That said, locomotion in the early time points of the NIL + cocaine test was higher than the NIL test, regardless of genotype, so the possibility that cocaine enhances exploratory behavior of RGS14 KO mice in a novel environment cannot be excluded.

A second explanation is that RGS14 KO mice are more neophobic and therefore hypoactive in the NIL test, but cocaine alleviates neophobia and increases locomotor behavior of RGS14 KO mice in a novel environment above the level of WT controls. This explanation also seems unlikely to account for the cocaine-induced reversal of NIL phenotypes; the dose of cocaine we administered is known to be anxiogenic (Blanchard and Blanchard 1999; Schank et al. 2008) and thus would be expected to decrease (not increase) overall NIL by enhancing neophobia.

The third explanation, which we endorse, is that RGS14 KO mice are more anxious in the NIL, CIL, and the NIL + cocaine tests. This interpretation accounts for the consistently augmented thigmotaxis behavior in the RGS14 KO mice, including the hthigmotaxis ratio in the NIL + cocaine test specific to RGS14 KO mice. Taken together, these findings suggest that high dose cocaine exacerbated neophobia (Blanchard and Blanchard 1999; Schank et al. 2008) in RGS14 KO, with thigmotaxis proving a more consistent metric than total locomotor activity.

### c-fos and pERK induction are increased in subregions of the hippocampus and amygdala of RGS14 KO mice following exposure to cocaine and novelty

We found that c-fos induction was intensely upregulated in CA1, CA2, and CeA of RGS14 KO mice compared to WT controls. We did not detect differences in c-fos induction between genotypes in Pir Ctx, subregions of the NAcc, or hippocampal area CA3. When we analyzed pERK expression in the same regions, an identical pattern emerged, with RGS14 KO mice demonstrating more pERK+ cells in CA1, CA2, and CeA, but not in Pir Ctx, NAcc, or CA3.

The remarkable correlation between c-fos and pERK induction following cocaine exposure is consistent with previous findings (Valjent et al. 2000; Valjent et al. 2006; Valjent et al. 2004). Pharmacological studies indicate that cocaine-induced hyperlocomotion and c-fos expression require DAergic signaling through D1Rs (Fricks-Gleason and Marshall 2011; Karlsson et al. 2008), glutamatergic signaling through NMDARs (Sun et al. 2008; Torres and Rivier 1993), and activation of the Ras/Raf/MEK/ERK pathway (Lu et al. 2006; Papale et al. 2016; Sun et al. 2016).

Exposure to novel environments induces expression of c-fos in dorsal hippocampal subfields CA1 and CA3, which are required for spatial learning and memory (Kempadoo et al. 2016; Moreno-Castilla et al. 2017; Wagatsuma et al. 2018). While CA2 is conventionally implicated in social recognition memory (Carstens and Dudek 2019; Dudek et al. 2016), the firing rate and place fields of CA2 pyramidal cells are modulated by novel environments (Alexander et al. 2016; Chen et al. 2020; Mankin et al. 2015), and chemogenetic manipulation of CA2 enhances freezing during fear learning (Alexander et al. 2019). Psychostimulants also induce c-fos and pERK in dorsal hipocampus, which is required for acquisition of context-associated psychostimulant behaviors (Koob and Volkow 2016; Kutlu and Gould 2016; Lu et al. 2006).

Novelty and psychostimulants reliably increase expression of pERK and c-fos within the CeA (Neisewander et al. 2000; Papa et al. 1993; Sanguedo et al. 2016; Valjent et al. 2004), a subregion of the amygdala implicated in stress-induced anxiety, aversive learning, and drug craving (Fadok et al. 2018; Lu et al. 2005). In a rodent model of anxiety induced by chronic pain, injury-associated increases in thigmotaxis were highly correlated with c-fos induction in the CeA (Morland et al. 2016). The relative increase of c-fos and ERK in the CeA of RGS14 KO mice compared to WT further support our interpretation that the augmented locomotor response of RGS14 KO mice in the NIL + cocaine test is attributable to anxiogenic effects of cocaine. While much is known about the role of the CeA in innate anxiety and drug responses (Fadok et al. 2018), the specific function of RGS14 within this region has not been studied.

### Proposed signaling model: RGS14 suppresses ERK signaling through interactions with Ras and possibly CaMKII

The unique tandem RBD region of RGS14 is a critical site for interactions with Ras and CaMKII (Evans et al. 2018a; Willard et al. 2009; Zhao et al. 2013). Increased intracellular Ca^2+^/calmodulin activates Ras and CaMKII, either of which can increase pERK and activate the transcription factor CREB to promote expression of IEGs (e.g., c-fos) and ultimately alter neuroplasticity. Although there is functional evidence that RGS14 sequesters Ca^2+^-activated Ras to suppress the Ras/Raf/MEK/ERK signaling axis (Shu et al. 2010), the physiological significance of RGS14 and CaMKII interaction has not been fully determined (Evans et al. 2018a). That said, the physiologically abnormal synaptic plasticity at CA3 → CA2 synapses in RGS14 KO mice requires NMDAR-dependent Ca^2+^/CaMKII signaling, in addition to PKA and pERK (Evans et al. 2018b; Lee et al. 2010). These findings suggests that RGS14 may suppress multiple signaling pathways related to neuronal excitability, Ca^2+^ signaling, and structural plasticity (Evans et al. 2018a; Li et al. 2016; Shu et al. 2010).

Given the striking overlap in the regional expression of RGS14 and striatal-enriched protein tyrosine phosphatase (STEP) (Boulanger et al. 1995; Venkitaramani et al. 2009), an attractive hypothesis of RGS14 function is that it acts similarly to STEP to regulate intracellular signaling events in neurons when glutamate and DA receptors are activated simultaneously (Braithwaite et al. 2006; Goebel-Goody et al. 2012; Nestler 2001; Russo et al. 2010). STEP inactivates pERK under basal conditions and is inactivated by PKA following D1R activation, thus disinhibiting pERK (Goebel-Goody et al. 2012). We propose that the NMDAR-dependent branch of the second messenger pathway could be similarly modulated by RGS14 (Fig. 6), which is not a phosphatase but reduces Ras/Raf/MEK/ERK signaling (Li et al. 2016). We also speculate that RGS14 may directly block Ca^2+^-dependent CaMKII signaling (Evans et al. 2018a), providing another molecular brake on Ca^2+^ and pERK signaling (Fig. 6).

**Figure 6.**
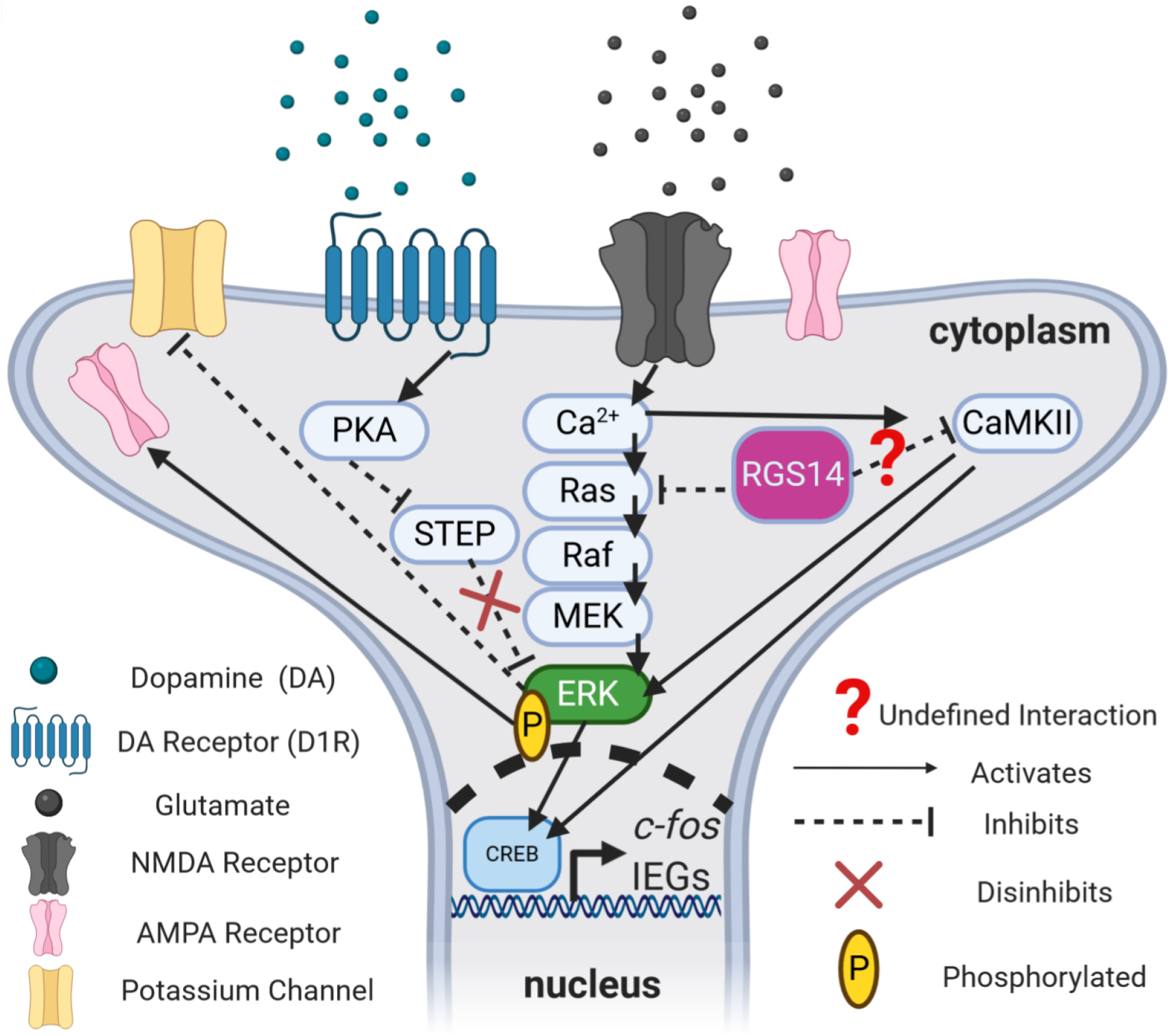
Proposed signaling model for RGS14-mediated suppression of cocaine-induced locomotion and gene expression. Phosphorylated ERK (pERK) is active and promotes rapid “cytoplasmic” effects by inactivating potassium efflux channels or increasing the insertion of ionotropic α-amino-3-hydroxy-5-methyl-4-isoxazolepropionic acid (AMPA) receptors for glutamate, as well as slower “nuclear” effects involving activation of the transcription factor CREB and subsequent upregulation of activity-dependent immediate early genes (IEGs) like *c-fos* that are involved in synaptic plasticity. The protein STEP dephosphorylates pERK and inactivates it, but STEP itself is inactivated following PKA activation downstream of D1 receptor (D1R) stimulation by dopamine (DA). Thus, D1R activation by DA disinhibits pERK by inhibiting the phosphatase activity of STEP towards pERK. Similarly, RGS14 may act to suppress pERK signaling by inhibiting Ca^2+^-dependent signaling downstream of NMDAR stimulation via known interactions with Ca^2+^-activated Ras and undefined interactions with activated CaMKII. These signaling interactions may occur either in the cytosol or in the nucleus, where RGS14 can shuttle. *Created with BioRender.com*.

### Limitations and future directions

A potential limitation of our study is that we cannot definitively attribute the locomotor-activating effects of cocaine in the RGS14 KOs to alterations in DA signaling. Cocaine increases synaptic concentrations of DA, NE, and 5-HT by inhibiting their respective transporters (DAT, NET, and SERT) (Schmidt and Weinshenker 2014). Moreover, activation of forebrain target regions of the monoamine modulatory systems following cocaine exposure also increases glutamatergic signaling within the mesocorticolimbic motivation circuitry (Baker et al. 2002). While prior work strongly suggests that CIL requires cocaine binding to DAT (Chen et al. 2006), additional locomotor experiments using the selective DAT antagonist GBR-12783 (Drouin et al. 2002) could further elucidate the role of DAT inhibition in the RGS14 KO cocaine-induced phenotypes reported here. Targeted pharmacological approaches, such as site-specific infusion of D1R and NMDAR antagonists into the limbic regions of RGS14 KO mice prior to systemic cocaine administration, could also reveal whether simultaneous DA and glutamate signaling is critical for enhanced locomotor responses to cocaine. Strategies to delete RGS14 in discrete brain regions would also be useful for determining the neuroanatomical substrates underlying the role of this protein in responses to novelty and psychostimulants.

Because the experiments in our study only assessed acute cocaine effects, future studies will be needed to investigate RGS14 function in the context of chronic cocaine exposure. Given the role of RGS14 in modulating plasticity, it would be particularly interesting to evaluate whether the RGS14 KO mice exhibit enhanced development and/or expression of locomotor sensitization, as well as place preference after repeated exposures to cocaine (Koob and Simon 2009; Rahman et al. 2003). It is also important to note that while we identified an increase in pERK and c-fos in discrete brain regions of RGS14 KO mice following the NIL + cocaine test, future experiments with brain-penetrant MEK inhibitors (Michalak et al. 2020; Valjent et al. 2000) will be required to determine whether a causal relationship exists between pERK/c-fos levels in RGS14-ir brain regions and locomotion. In follow-up studies, it would also be interesting to correlate individual differences in CIL or NIL behavior (including thigmotaxis) with regional changes in pERK and cfos expression to refine our understanding of the relative contributions of these regions to specific behaviors (Morland et al. 2016).

### Conclusions

We identified a deficit in NIL and an enhancement in high-dose CIL in RGS14 KO mice that suggests a previously unappreciated sensitivity to novelty- and psychostimulant-induced anxiety in these mutant mice (Paine et al. 2002; Pawlak et al. 2008). Although previous behavioral assessments of RGS14 KO mice did not identify an anxiety-like phenotype (Lee et al. 2010), we found that RGS14 KO mice exhibit enhanced thigmotaxis behavior compared to WT under conditions of environmental novelty or high-dose cocaine, and that when combined these factors exerted an additive effect on thigmotaxis in RGS14 KO mice. Importantly, our findings illuminate previously unknown functions for RGS14 in the regulation of innate and drug-induced anxiety and locomotor behavior (Cryan and Sweeney 2011; Paine et al. 2002), which are biologically distinct from learning and memory (Alexander et al. 2019; Lee et al. 2010).

Finally, we observed marked increases in c-fos and pERK induction in subregions of the hippocampus and amygdala in RGS14 KO mice following exposure to high-dose cocaine and environmental novelty. We have also integrated our findings with those from previous studies to propose a new signaling model for RGS14 as a STEP-like regulator of Ca^2+^-dependent second messenger cascades in DA-receptive neurons (Fig. 6), which is fertile ground for future mechanistic studies of RGS14 function in cell culture models and *ex vivo* brain slice preparations.

## Acknowledgments

We thank C. Strauss for helpful editing of the manuscript.

